# Expanding a Database-derived Biomedical Knowledge Graph via Multi-relation Extraction from Biomedical Abstracts

**DOI:** 10.1101/730085

**Authors:** David N. Nicholson, Daniel S. Himmelstein, Casey S. Greene

## Abstract

Knowledge graphs support multiple research efforts by providing contextual information for biomedical entities, constructing networks, and supporting the interpretation of high-throughput analyses. These databases are populated via some form of manual curation, which is difficult to scale in the context of an increasing publication rate. Data programming is a paradigm that circumvents this arduous manual process by combining databases with simple rules and heuristics written as label functions, which are programs designed to automatically annotate textual data. Unfortunately, writing a useful label function requires substantial error analysis and is a nontrivial task that takes multiple days per function. This makes populating a knowledge graph with multiple nodes and edge types practically infeasible. We sought to accelerate the label function creation process by evaluating the extent to which label functions could be re-used across multiple edge types. We used a subset of an existing knowledge graph centered on disease, compound, and gene entities to evaluate label function re-use. We determined the best label function combination by comparing a baseline database-only model with the same model but added edge-specific or edge-mismatch label functions. We confirmed that adding additional edge-specific rather than edge-mismatch label functions often improves text annotation and shows that this approach can incorporate novel edges into our source knowledge graph. We expect that continued development of this strategy has the potential to swiftly populate knowledge graphs with new discoveries, ensuring that these resources include cutting-edge results.

## Introduction

Knowledge bases are important resources that hold complex structured and unstructured information. These resources have been used in important tasks such as network analysis for drug repurposing discovery [1,2,3] or as a source of training labels for text mining systems [4,5,6]. Populating knowledge bases often requires highly trained scientists to read biomedical literature and summarize the results [7]. This time-consuming process is referred to as manual curation. In 2007, researchers estimated that filling a knowledge base via manual curation would require approximately 8.4 years to complete [8]. The rate of publications continues to exponentially increase [9], so using only manual curation to fully populate a knowledge base has become impractical.

Relationship extraction has been studied as a solution towards handling the challenge posed by an exponentially growing body of literature [7]. This process consists of creating an expert system to automatically scan, detect and extract relationships from textual sources. Typically, these systems utilize machine learning techniques that require extensive corpora of well-labeled training data. These corpora are difficult to obtain, because they are constructed via extensive manual curation pipelines.

Distant supervision is a technique also designed to sidestep the dependence on manual curation and quickly generate large training datasets. This technique assumes that positive examples established in selected databases can be applied to any sentence that contains them [4]. The central problem with this technique is that generated labels are often of low quality which results in an expansive amount of false positives [10].

Ratner et al. [11] recently introduced “data programming” as a solution. Data programming is a paradigm that combines distant supervision with simple rules and heuristics written as small programs called label functions. These label functions are consolidated via a noise aware generative model that is designed to produce training labels for large datasets. Using this paradigm can dramatically reduce the time required to obtain sufficient training data; however, writing a useful label function requires a significant amount of time and error analysis. This dependency makes constructing a knowledge base with a myriad of heterogenous relationships nearly impossible as tens or possibly hundreds of label functions are required per relationship type.

In this paper, we seek to accelerate the label function creation process by measuring the extent to which label functions can be re-used across different relationship types. We hypothesize that sentences describing one relationship type may share linguistic features such as keywords or sentence structure with sentences describing other relationship types. We conducted a series of experiments to determine the degree to which label function re-use enhanced performance over distant supervision alone. We focus on relationships that indicate similar types of physical interactions (i.e., gene-binds-gene and compound-binds-gene) as well as different types (i.e., disease-associates-gene and compound-treats-disease). Re-using label functions could dramatically reduce the time required to populate a knowledge base with a multitude of heterogeneous relationships.

### Related Work

Relationship extraction is the process of detecting semantic relationships from a collection of text. This process can be broken down into three different categories: (1) the use of natural language processing techniques such as manually crafted rules and heuristics for relationship extraction (Rule Based Extractors), (2) the use of unsupervised methods such as co-occurrence scores or clustering to find patterns within sentences and documents (Unsupervised Extractors), and (3) the use of supervised or semi-supervised machine learning for classifying the presence of a relation within documents or sentences (Supervised Extractors). In this section, we briefly discuss selected efforts under each category.

### Rule Based Extractors

Rule based extractors rely heavily on expert knowledge to perform extraction. Typically, these systems use linguistic rules and heuristics to identify key sentences or phrases. For example, a hypothetical extractor focused on protein phosphorylation events would identify sentences containing the phrase “gene X phosphorylates gene Y” [12]. This phrase is a straightforward indication that two genes have a fundamental role in protein phosphorylation. Other phrase extractors have been used to identify drug-disease treatments [13], pharmcogenomic events [14] and protein-protein interactions [15,16]. These extractors provide a simple and effective way to extract sentences; however, they depend on extensive knowledge about the text to be properly constructed.

A sentence’s grammatical structure can also support relationship extraction via dependency trees. Dependency trees are data structures that depict a sentence’s grammatical relation structure in the form of nodes and edges. Nodes represent words and edges represent the dependency type each word shares between one another. For example, a possible extractor would classify sentences as a positive if a sentence contained the following dependency tree path: “gene X (subject)-> promotes (verb)<- cell death (direct object) <- in (preposition) <-tumors (object of preposition)” [17]. This approach provides extremely precise results, but the quantity of positive results remains modest as sentences appear in distinct forms and structure. Because of this limitation, recent approaches have incorporated methods on top of rule based extractors such as co-occurrence and machine learning systems [18,19]. We discuss the pros and cons of added methods in a later section. For this project, we constructed our label functions without the aid of these works; however, approaches discussed in this section provide substantial inspiration for novel label functions in future endeavors.

### Unsupervised Extractors

Unsupervised extractors detect relationships without the need of annotated text. Notable approaches exploit the fact that two entities can occur together in text. This event is referred to as co-occurrence. Extractors utilize these events by generating statistics on the frequency of entity pairs occurring in text. For example, a possible extractor would say gene X is associated with disease Y, because gene X and disease Y appear together more often than individually [20]. This approach has been used to establish the following relationship types: disease-gene relationships [20,21,22,23,24,25], protein-protein interactions [24,26,27], drug-disease treatments [28], and tissue-gene relations [29]. Extractors using the co-occurrence strategy provide exceptional recall results; however, these methods may fail to detect underreported relationships, because they depend on entity-pair frequency for detection. Junge et al. created a hybrid approach to account for this issue using distant supervision to train a classifier to learn the context of each sentence [30]. Once the classifier was trained, they scored every sentence within their corpus, and each sentence’s score was incorporated into calculating co-occurrence frequencies to establish relationship existence [30]. Co-occurrence approaches are powerful in establishing edges on the global scale; however, they cannot identify individual sentences without the need for supervised methods.

Clustering is an unsupervised approach that extracts relationships from text by grouping similar sentences together. Percha et al. used this technique to group sentences based on their grammatical structure [31]. Using Stanford’s Core NLP Parser [32], a dependency tree was generated for every sentence in each Pubmed abstract [31]. Each tree was clustered based on similarity and each cluster was manually annotated to determine which relationship each group represented [31]. For our project we incorporated the results of this work as domain heuristic label functions. Overall, unsupervised approaches are desirable since they do not require well-annotated training data. Such approaches provide excellent recall; however, performance can be limited in terms of precision when compared to supervised machine learning methods [33,34].

### Supervised Extractors

Supervised extractors consist of training a machine learning classifier to predict the existence of a relationship within text. These classifiers require access to well-annotated datasets, which are usually created via some form of manual curation. Previous work consists of research experts curating their own datasets to train classifiers [35,36,37,38,39]; however, there have been community-wide efforts to create datasets for shared tasks [40,41,42]. Shared tasks are open challenges that aim to build the best classifier for natural language processing tasks such as named entity tagging or relationship extraction. A notable example is the BioCreative community that hosted a number of shared tasks such as predicting compound-protein interactions (BioCreative VI track 5) [41] and compound induced diseases [42]. Often these datasets are well annotated, but are modest in size (2,432 abstracts for BioCreative VI [41] and 1500 abstracts for BioCreative V [42]). As machine learning classifiers become increasingly complex, these small dataset sizes cannot suffice. Plus, these multitude of datasets are uniquely annotated which can generate noticeable differences in terms of classifier performance [42]. Overall, obtaining large well-annotated datasets still remains as an open non-trivial task.

Before the rise of deep learning, a classifier that was most frequently used was support vector machines. This classifier uses a projection function called a kernel to map data onto a high dimensional space so datapoints can be easily discerned between classes [43]. This method was used to extract disease-gene associations [35,44,45], protein-protein interactions[19,46,47] and protein docking information [48]. Generally, support vector machines perform well on small datasets with large feature spaces but are slow to train as the number of datapoints becomes asymptotically large.

Deep learning has been increasingly popular as these methods can outperform common machine learning methods [49]. Approaches in this field consist of using various neural network architectures, such as recurrent neural networks [50,51,52,53,54,55] and convolutional neural networks [51,54,56,57,58], to extract relationships from text. In fact approaches in this field were the winning model within the BioCreative VI shared task [41,59]. Despite the substantial success of these models, they often require large amounts of data to perform well. Obtaining large datasets is a time-consuming task, which makes training these models a non-trivial challenge. Distant supervision has been used as a solution to fix the barren amount of large datasets [4]. Approaches have used this paradigm to extract chemical-gene interactions [54], disease-gene associations [30] and protein-protein interactions [30,54,60]. In fact, efforts done in [60] served as one of the motivating rationales for our work.

Overall, deep learning has provided exceptional results in terms of relationships extraction. Thus, we decided to use a deep neural network as our discriminative model.

## Methods and Materials

### Hetionet

Hetionet v1 [3] is a large heterogenous network that contains pharmacological and biological information. This network depicts information in the form of nodes and edges of different types: nodes that represent biological and pharmacological entities and edges which represent relationships between entities. Hetionet v1 contains 47,031 nodes with 11 different data types and 2,250,197 edges that represent 24 different relationship types (Figure 1). Edges in Hetionet v1 were obtained from open databases, such as the GWAS Catalog [61] and DrugBank [62]. For this project, we analyzed performance over a subset of the Hetionet v1 edge types: disease associates with a gene (DaG), compound binds to a gene (CbG), compound treating a disease (CtD) and gene interacts with gene (GiG) (bolded in Figure 1).

**Figure 1:**
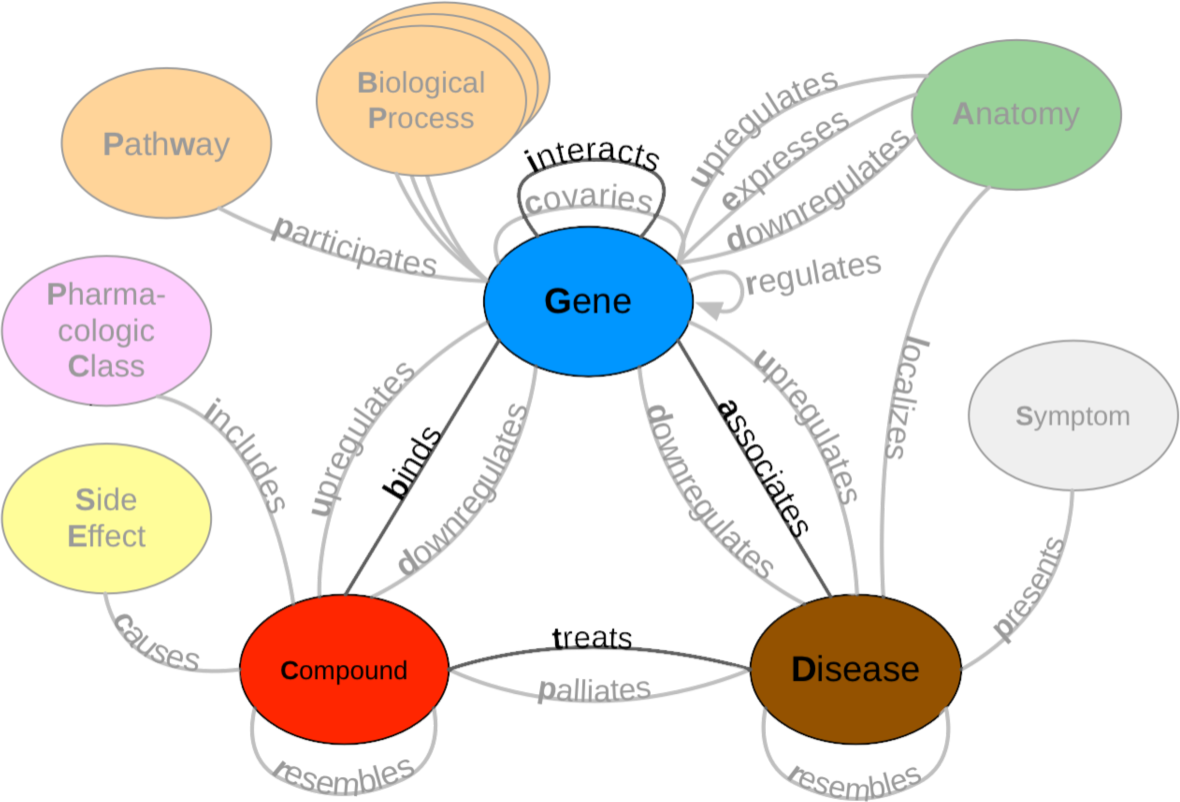
A metagraph (schema) of Hetionet v1 where biomedical entities are represented as nodes and the relationships between them are represented as edges. We examined performance on the highlighted subgraph; however, the long-term vision is to capture edges for the entire graph.

### Dataset

We used PubTator [63] as input to our analysis. PubTator provides MEDLINE abstracts that have been annotated with well-established entity recognition tools including DNorm [64] for disease mentions, GeneTUKit [65] for gene mentions, Gnorm [66] for gene normalizations and a dictionary based search system for compound mentions [67]. We downloaded PubTator on June 30, 2017, at which point it contained 10,775,748 abstracts. Then we filtered out mention tags that were not contained in Hetionet v1. We used the Stanford CoreNLP parser [32] to tag parts of speech and generate dependency trees. We extracted sentences with two or more mentions, termed candidate sentences. Each candidate sentence was stratified by co-mention pair to produce a training set, tuning set and a testing set (shown in Supplemental Table 2). Each unique co-mention pair was sorted into four categories: (1) in Hetionet v1 and has sentences, (2) in Hetionet v1 and doesn’t have sentences, (3) not in Hetionet v1 and does have sentences and (4) not in Hetionet v1 and doesn’t have sentences. Within these four categories each pair is randomly assigned their own individual partition rank (a continuous number between 0 and 1). Any rank lower than 0.7 is sorted into the training set, while any rank greater than 0.7 and lower than 0.9 is assigned to the tuning set. The rest of the pairs with a rank greater than or equal to 0.9 is assigned to the test set. Sentences that contain more than one co-mention pair are treated as multiple individual candidates. We hand labeled five hundred to a thousand candidate sentences of each edge type to obtain a ground truth set (Supplemental Table 2)^1^.

### Label Functions for Annotating Sentences

The challenge of having too few ground truth annotations is common to many natural language processing settings, even when unannotated text is abundant. Data programming circumvents this issue by quickly annotating large datasets by using multiple noisy signals emitted by label functions [11]. Label functions are simple pythonic functions that emit: a positive label (1), a negative label (−1) or abstain from emitting a label (0). These functions can be grouped into multiple categories (see Supplement Methods). We combined these functions using a generative model to output a single annotation, which is a consensus probability score bounded between 0 (low chance of mentioning a relationship) and 1 (high chance of mentioning a relationship). We used these annotations to train a discriminative model that makes the final classification step.

### Experimental Design

Being able to re-use label functions across edge types would substantially reduce the number of label functions required to extract multiple relationships from biomedical literature. We first established a baseline by training a generative model using only distant supervision label functions designed for the target edge type (see Supplemental Methods). For example, in the Gene interacts Gene (GiG) edge type we used label functions that returned a 1 if the pair of genes were included in the Human Interaction database [68], the iRefIndex database [69] or in the Incomplete Interactome database [70]. Then we compared the baseline model with models that also included text and domain-heuristic label functions. Using a sampling with replacement approach, we sampled these text and domain-heuristic label functions separately within edge types, across edge types, and from a pool of all label functions. We compared within-edge-type performance to across-edge-type and all-edge-type performance. For each edge type we sampled a fixed number of label functions consisting of five evenly spaced numbers between one and the total number of possible label functions. We repeated this sampling process 50 times for each point. Furthermore, at each point we also trained the discriminative model using annotations from the generative model trained on edge-specific label functions (see Supplemental Methods). We report performance of both models in terms of the area under the receiver operating characteristic curve (AUROC) and the area under the precision-recall curve (AUPR). Ensuing model evaluations, we quantified the number of edges we could incorporate into Hetionet v1. Using a calibrated discriminative model (see Supplemental Methods), we scored every candidate sentence within our dataset and grouped candidates based on their mention pair. We took the max score within each candidate group and this score represents the probability of the existence of an edge. We established edges by using a cutoff score that produced an equal error rate between the false positives and false negatives. We report the number of preexisting edges we could recall as well as the number of novel edges we can incorporate. Lastly, we compared our framework with a previously established unsupervised approach [30].

## Results

### Generative Model Using Randomly Sampled Label Functions

Creating label functions is a labor-intensive process that can take days to accomplish. We sought to accelerate this process by measuring the extent to which label functions can be reused. Our hypothesis was that certain edge types share similar linguistic features such as keywords and/or sentence structure. This shared characteristic would make certain edge types amenable to label function reuse. We designed a set of experiments to test this hypothesis on an individual level (edge vs edge) as well as a global level (collective pool of sources). We observed that performance increased when edge-specific label functions were added to an edge-specific baseline model, while label function reuse usually provided less benefit (AUROC Figure 2, AUPR Supplemental Figure 5). We also evaluated randomly selecting label functions from among all sets and observed similar performance (AUROC Supplemental Figure 6, AUPR Supplemental Figure 7) The quintessential example of this overarching trend is the Compound treats Disease (CtD) edge type, where edge-specific label functions always outperformed transferred label functions. However, there are hints of label function transferability for selected edge types and label function sources. Performance increases as more CbG label functions are incorporated to the GiG baseline model and vice versa. This suggests that sentences for GiG and CbG may share similar linguistic features or terminology that allows for label functions to be reused. Perplexingly, edge-specific Disease associates Gene (DaG) label functions did not improve performance over label functions drawn from other edge types. Overall, only CbG and GiG showed significant signs of reusability which suggests label functions could be shared between the two edge types.

**Figure 2:**
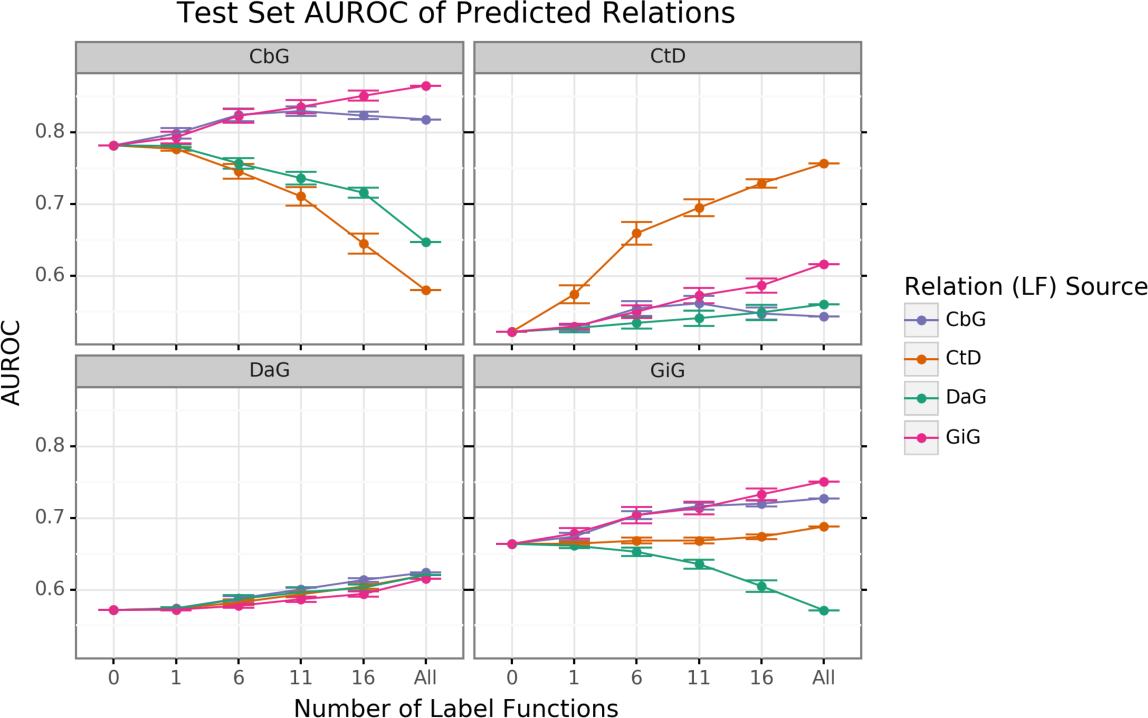
Edge-specific label functions are better performing than edge-mismatch label functions, but certain mismatch situations show signs of successful transfer. Each line plot header depicts the edge type the generative model is trying to predict, while the colors represent the source of label functions. For example, orange represents sampling label functions designed to predict the Compound treats Disease (CtD) edge type. The x axis shows the number of randomly sampled label functions being incorporated into the database-only baseline model (point at 0). The y axis shows area under the receiver operating curve (AUROC). Each point on the plot shows the average of 50 sample runs, while the error bars show the 95% confidence intervals of all runs. The baseline and “All” data points consist of sampling from the entire fixed set of label functions.

We found that sampling from all label function sources at once usually underperformed relative to edge-specific label functions (Supplemental Figures 6 and 7). As more label functions were sampled, the gap between edge-specific sources and all sources widened. CbG is a prime example of this trend (Supplemental Figures 6 and 7), while CtD and GiG show a similar but milder trend. DaG was the exception to the general rule: the pooled set of label functions improved performance over the edge-specific ones, which aligns with the previously observed results for individual edge types (Figure 2). The decreasing trend when pooling all label functions supports the notion that label functions cannot easily transfer between edge types (exception being CbG on GiG and vice versa).

### Discriminative Model Performance

The discriminative model is designed to augment performance over the generative model by incorporating textual features along with estimated training labels. The discriminative model is a piecewise convolutional neural network trained over word embeddings (See Methods and Materials). We found that the discriminative model generally out-performed the generative model as more edge-specific label functions are incorporated (Figure 3 and Supplemental Figure 8). The discriminative model’s performance is often poorest when very few edge-specific label functions are added to the baseline model (seen in Disease associates Gene (DaG), Compound binds Gene (CbG) and Gene interacts Gene (GiG)). This suggests that generative models trained with more label functions produce outputs that are more suitable for training discriminative models. An exception to this trend is Compound treats Disease (CtD) where the discriminative model out-performs the generative model at all levels of sampling. We observed the opposite trend with the Compound-binds-Gene (CbG) edges: the discriminative model was always poorer or indistinguishable from the generative model. Interestingly, the AUPR for CbG plateaus below the generative model and decreases when all edge-specific label functions are used (Supplemental Figure 8). This suggests that the discriminative model might be predicting more false positives in this setting. Incorporating more edge-specific label functions usually improves performance for the discriminative model over the generative model.

**Figure 3:**
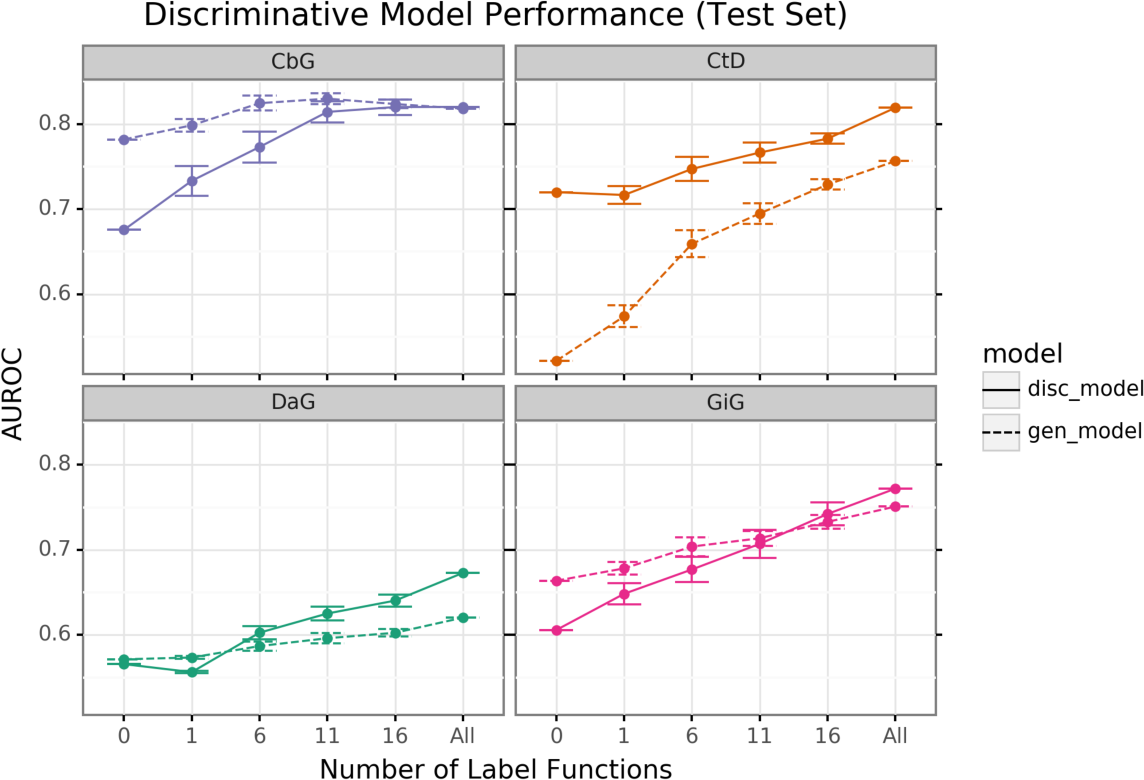
The discriminative model usually improves at a faster rate than the generative model as more edge-specific label function are included. The line plot headers represent the specific edge type the discriminative model is trying to predict. The x-axis shows the number of randomly sampled label functions that are incorporated into the baseline model (point at 0). The y axis shows the area under the receiver operating curve (AUROC). Each datapoint represents the average of 50 sample runs and the error bars represent the 95% confidence interval of each run. The baseline and “All” data points consist of sampling from the entire fixed set of label functions.

## Discussion

We measured the extent to which label functions can be re-used across multiple edge types to extract relationships from literature. Through our sampling experiment, we found that adding edge-specific label functions increases performance for the generative model (Figure 2). We found that label functions designed from relatively related edge types can increase performance (Gene interacts Gene (GiG) label functions predicting the Compound binds Gene (CbG) edge and vice versa), while the Disease associates Gene (DaG) edge type remained agnostic to label function sources (Figure 2 and Supplemental Figure 5). Furthermore, we found that using all label functions at once generally hurts performance with the exception being the DaG edge type (Supplemental Figures 6 and 7). One possibility for this observation is that DaG is a broadly defined edge type. For example, DaG may contain many concepts related to other edge types such as Disease (up/down) regulating a Gene, which makes it more agnostic to label function sources (examples highlighted in our annotated sentences).

Regarding the discriminative model, adding edge-specific label function substantially improved performance for two out of the four edge types (Compound treats Disease (CtD) and Disease associates Gene (DaG)) (Figure 3 and Supplemental Figure 8). Gene interacts Gene (GiG) and Compound binds Gene (CbG) discriminative models showed minor improvements compared to the generative model, but only when nearly all edge-specific label functions are included (Figure 3 and Supplemental Figure 8). We came across a large amount of spurious gene mentions when working with the discriminative model and believe that these mentions contributed to CbG and GiG’s hindered performance. We encountered difficulty in calibrating each discriminative model (Supplemental Figure 9). The temperature scaling algorithm appears to improve calibration for the highest scores for each model but did not successfully calibrate throughout the entire range of predictions. Improving performance for all predictions may require more labeled examples or may be a limitation of the approach in this setting. Even with these limitations, this early-stage approach could recall many existing edges from an existing knowledge base, Hetionet v1, and suggest many new high-confidence edges for inclusion (Supplemental Figure 10). Our findings suggest that further work, including an expansion of edge types and a move to full text from abstracts, may make this approach suitable for building continuously updated knowledge bases to address drug repositioning and other biomedical challenges.

## Conclusion and Future Direction

Filling out knowledge bases via manual curation can be an arduous and erroneous task [8]. As the rate of publications increases, relying on manual curation alone becomes impractical. Data programming, a paradigm that uses label functions as a means to speed up the annotation process, can be used as a solution for this problem. An obstacle for this paradigm, however, is creating useful label functions, which takes a considerable amount of time. We tested the feasibility of reusing label functions as a way to reduce the total number of label functions required for strong prediction performance. We conclude that label functions may be re-used with closely related edge types, but that re-use does not improve performance for most pairings. The discriminative model’s performance improves as more edge-specific label functions are incorporated into the generative model; however, we did notice that performance greatly depends on the annotations provided by the generative model.

This work sets up the foundation for creating a common framework that mines text to create edges. Within this framework we would continuously incorporate new knowledge as novel findings are published, while providing a single confidence score for an edge via sentence score consolidation. As opposed to many existing knowledge graphs (for example, Hetionet v1 where text-derived edges generally cannot be exactly attributed to excerpts from literature [3,71]), our approach has the potential to annotate each edge based on its source sentences. In addition, edges generated with this approach would be unencumbered from upstream licensing or copyright restrictions, enabling openly licensed hetnets at a scale not previously possible [72,73,74]. New multitask learning [75] strategies may make it even more practical to reuse label functions to construct continuously updating literature-derived knowledge graphs.

## Supplemental Information

An online version of this manuscript is available at https://greenelab.github.io/text_mined_hetnet_manuscript/. Source code for this work is available under open licenses at: https://github.com/greenelab/snorkeling/.

## Acknowledgements

The authors would like to thank Christopher Ré’s group at Stanford University, especially Alex Ratner and Steven Bach, for their assistance with this project. We also want to thank Graciela Gonzalez-Hernandez for her advice and input with this project. This work was support by Grant GBMF4552 from the Gordon Betty Moore Foundation.

## Footnotes

1 Labeled sentences are available here.

