## Supplemental Materials for "Expanding a Database-derived Biomedical Knowledge Graph via Multi-relation Extraction from Biomedical Abstracts"

### Supplemental Methods

#### Label Function Categories

Label functions can be constructed in a multitude of ways; however, many label functions share similar characteristics with one another. We grouped these characteristics into the following categories: databases, text patterns and domain heuristics. Most of our label functions fall into the text pattern category, while the others were distributed across the database and domain heuristic categories (Table 1). Further, we described each category and provided an example that refers to the following candidate sentence: "PTK6 may be a novel therapeutic target for pancreatic cancer".

**Databases**: These label functions incorporate existing databases to generate a signal, as seen in distant supervision [4]. These functions detect if a candidate sentence's co-mention pair is present in a given database. If the pair is present, our label function emits a positive label and abstains otherwise. If the pair is not present in any existing database, a separate label function emits a negative label. We used a separate label function to prevent a label imbalance problem that we encountered during development: emitting positives and negatives from the same label function causes downstream classifiers to generate almost exclusively negative predictions.

$$\Lambda_{DB}(D, G) = \begin{cases} 1 & (D, G) \in DB \\ 0 & otherwise \end{cases}$$

$$\Lambda_{\neg DB}(D, G) = \begin{cases} -1 & (D, G) \notin DB \\ 0 & otherwise \end{cases}$$

**Domain Heuristics**: These label functions used results from published text-based analyses to generate a signal. For our project, we used dependency path cluster themes generated by Percha et al. [31]. If a candidate sentence's dependency path belonged to a previously generated cluster, then the label function emitted a positive label and abstained otherwise.

$$\Lambda_{DH}(D, G) = \begin{cases} 1 & Candidate\ Sentence \in Cluster\ Theme \\ 0 & otherwise \end{cases}$$

**Text Patterns**: These label functions are designed to use keywords and sentence context to generate a signal. For example, a label function could focus on the number of words between two mentions or focus on the grammatical structure of a sentence. These functions emit a positive or negative label depending on the context.

$$\Lambda_{TP}(D, G) = \begin{cases} 1 & ''\ target\ '' \in Candidate\ Sentence \\ 0 & otherwise \end{cases}$$

$$\Lambda_{TP}(D, G) = \begin{cases} -1 & ''\ VB\ '' \notin pos\_tags(Candidate\ Sentence) \\ 0 & otherwise \end{cases}$$

Each text pattern label function was constructed by manual examination of sentences within the training set. For example, in the candidate sentence above, one would extract the keywords "novel therapeutic target" and incorporate them in a text pattern label function. After initial construction, we tested and augmented the label function using sentences in the tune set. We repeated this process for every label function in our repertoire.

**Table 1:** The distribution of each label function per relationship.

| Relationship | Databases (DB) | Text Patterns (TP) | Domain Heuristics (DH) |
|---|---|---|---|
| DaG | 7 | 20 | 10 |
| CtD | 3 | 15 | 7 |
| CbG | 9 | 13 | 7 |
| GiG | 9 | 20 | 8 |

### Training Models

#### Generative Model

The generative model is a core part of this automatic annotation framework. It integrates multiple signals emitted by label functions and assigns a training class to each candidate sentence. This model assigns training classes by estimating the joint probability distribution of the latent true class ($Y$) and label function signals ($\Lambda$), ($P_\theta(\Lambda, Y)$). Assuming each label function is conditionally independent, the joint distribution is defined as follows:

$$P_\theta(\Lambda, Y) = \frac{\exp(\sum_{i=1}^{m} \theta^T F_i(\Lambda, y))}{\sum_{\Lambda'} \sum_{y'} \exp(\sum_{i=1}^{m} \theta^T F_i(\Lambda', y'))}$$

where $m$ is the number of candidate sentences, $F$ is the vector of summary statistics and $\theta$ is a vector of weights for each summary statistic. The summary statistics used by the generative model are as follows:

$$F_{i,j}^{Lab}(\Lambda, Y) = \mathbb{1}\{\Lambda_{i,j} \neq 0\}$$
$$F_{i,j}^{Acc}(\Lambda, Y) = \mathbb{1}\{\Lambda_{i,j} = y_{i,j}\}$$

*Lab* is the label function's propensity (the frequency of a label function emitting a signal). *Acc* is the individual label function's accuracy given the training class. This model optimizes the weights ($\theta$) by minimizing the negative log likelihood:

$$\hat{\theta} = argmin_\theta - \sum_{\Lambda} \sum_{Y} log P_\theta(\Lambda, Y)$$

In the framework we used predictions from the generative model, $\hat{Y} = P_{\hat{\theta}}(Y \mid \Lambda)$, as training classes for our dataset [75,76].

#### Discriminative Model

The discriminative model is a neural network trained to produce classification labels by integrating predicted probabilities from the generative model along with sentence representations via word embeddings. The goal of this combined approach is to develop models that learn text features associated with the overall task, beyond the supplied label functions. We used a piecewise convolutional neural network that contains multiple kernel filters as our discriminative model. We built a network with multiple filters using a fixed width of 300 (size of word embeddings) and a fixed height of 7 (Figure 4). We chose a fixed height of 7 because this height was previously reported to optimize performance in relationship classification [77]. We trained this model for 15 epochs using the Adam optimizer [78] with PyTorch's default parameter settings and a learning rate of 0.001 that decreases by half every epoch until the lower bound of 1e-5 is reached, which we observed was often sufficient for convergence. We added a L2 penalty (lambda=0.002) on the network weights to prevent overfitting. Lastly, we added a dropout layer (p=0.25) between the fully connected layer and the softmax layer.

**Figure 4:** The architecture of the discriminative model was a convolutional neural network. We performed a convolution step using multiple filters. The filters generated a feature map that was sent into a maximum pooling layer that was designed to extract the largest feature in each map. The extracted features were concatenated into a singular vector that was passed into a fully connected network. The fully connected network had 300 neurons for the first layer, 100 neurons for the second layer and 50 neurons for the last layer. The last step of the fully connected network was to generate predictions using a softmax layer.

#### Word Embeddings

Word embeddings are representations that map individual words to real valued vectors of user-specified dimensions. These embeddings have been shown to capture the semantic and syntactic information between words [79]. We trained Facebook's fastText [80] using all candidate sentences for each individual relationship pair to generate word embeddings. FastText uses a skip-gram model [81] that aims to predict the surrounding context for a candidate word and pairs the model with a novel scoring function that treats each word as a bag of character n-grams. We trained this model for 20 epochs using a window size of 2 and generated 300-dimensional word embeddings. We use the optimized word embeddings as input to our discriminative model.

#### Calibration of the Discriminative Model

Often many tasks require a machine learning model to output reliable probability predictions. A model is well calibrated if the probabilities emitted from the model match the observed probabilities. For example, a well-calibrated model that assigns a class label with 80% probability should have that class appear 80% of the time. Deep neural network models can often be poorly calibrated [82,83]. These models are usually over-confident in their predictions. For this reason, we calibrated our convolutional neural network using temperature scaling [82]. Temperature scaling uses a parameter T to scale each value of the logit vector (z) before being passed into the softmax (SM) function.

$$\sigma_{SM}(\frac{z_i}{T}) = \frac{\exp(\frac{z_i}{T})}{\sum_i \exp(\frac{z_i}{T})}$$

We found the optimal T by minimizing the negative log likelihood (NLL) of the tune set.

### Supplemental Figures

#### Generative Model Using Randomly Sampled Label Functions

#### Individual Sources

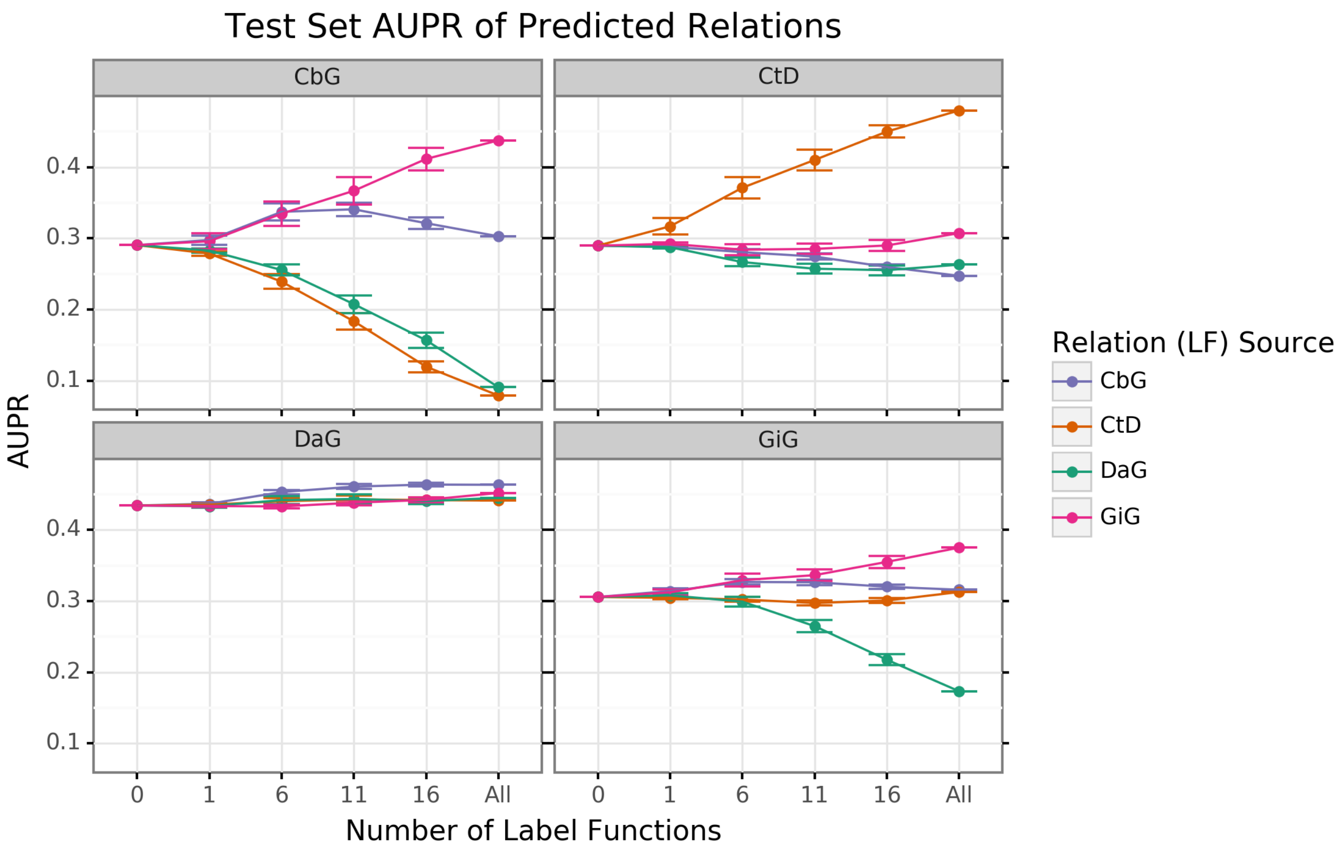


**Figure 5:** Edge-specific label functions improves performance over edge-mismatch label functions. Each line plot header depicts the edge type the generative model is trying to predict, while the colors represent the source of label functions. For example orange represents sampling label functions designed to predict the Compound treats Disease (CtD) edge type. The x axis shows the number of randomly sampled label functions being incorporated into the database-only baseline model (point at 0). The y axis shows area under the precision recall curve (AUPR). Each point on the plot shows the average of 50 sample runs, while the error bars show the 95% confidence intervals of all runs. The baseline and "All" data points consist of sampling from the entire fixed set of label functions.

#### Collective Pool of Sources

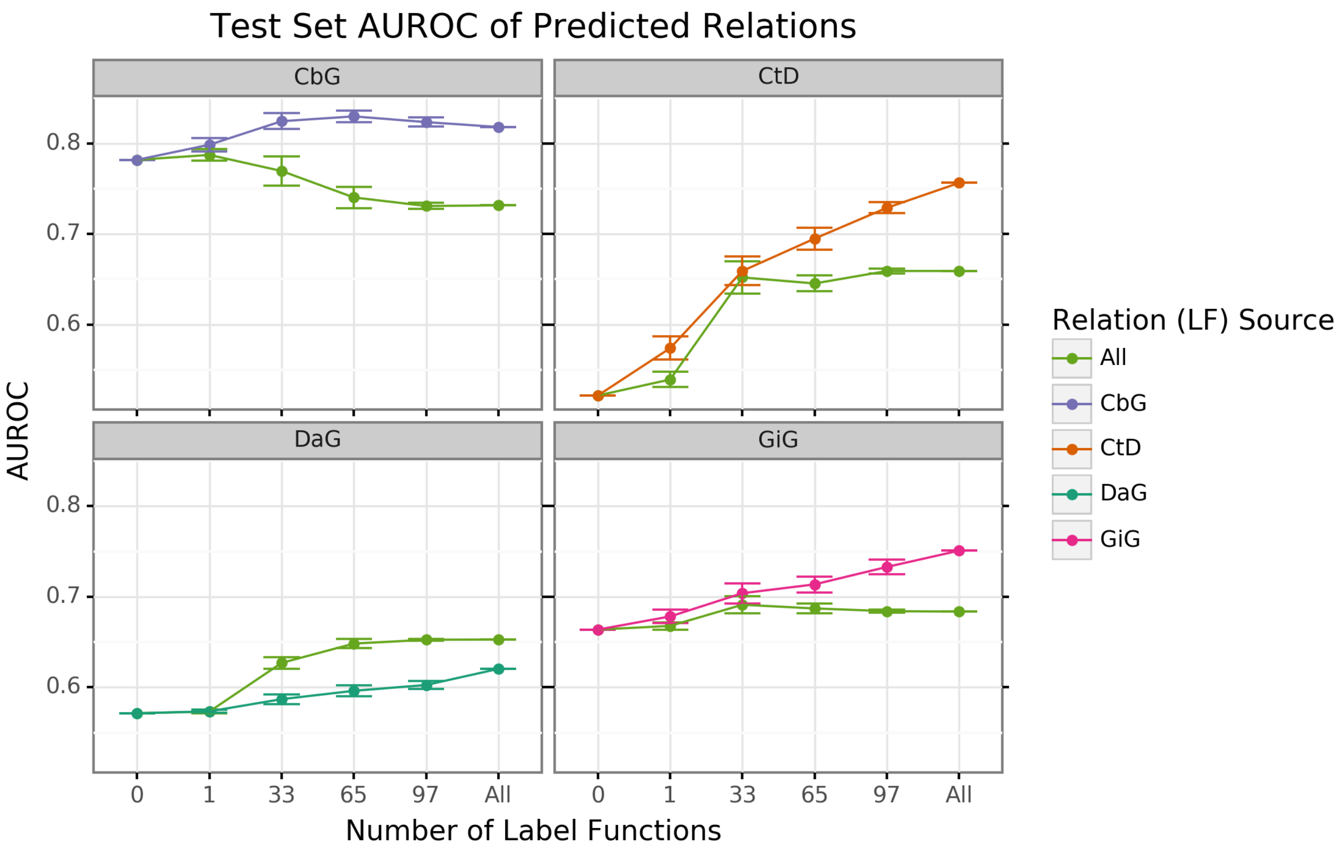


**Figure 6:** Using all label functions generally hinders generative model performance. Each line plot header depicts the edge type the generative model is trying to predict, while the colors represent the source of label functions. For example, orange represents sampling label functions designed to predict the Compound treats Disease (CtD) edge type. The x axis shows the number of randomly sampled label functions being incorporated into the database-only baseline model (point at 0). The y axis shows area under the receiver operating curve (AUROC). Each point on the plot shows the average of 50 sample runs, while the error bars show the 95% confidence intervals of all runs. The baseline and "All" data points consist of sampling from the entire fixed set of label functions.

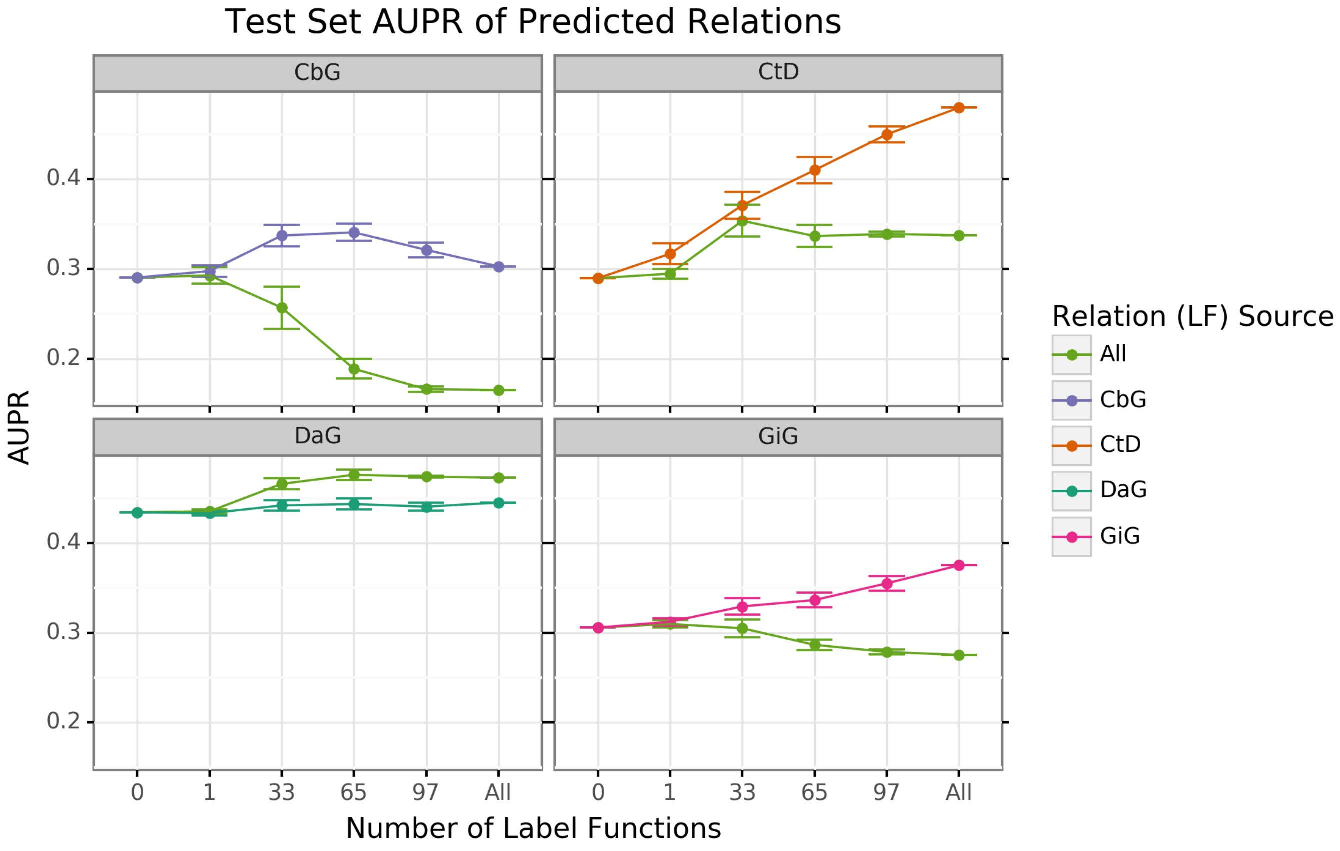

**Figure 7:** Using all label functions generally hinders generative model performance. Each line plot header depicts the edge type the generative model is trying to predict, while the colors represent the source of label functions. For example, orange represents sampling label functions designed to predict the Compound treats Disease (CtD) edge type. The x axis shows the number of randomly sampled label functions being incorporated into the database-only baseline model (point at 0). The y axis shows area under the precision recall curve (AUPR). Each point on the plot shows the average of 50 sample runs, while the error bars show the 95% confidence intervals of all runs. The baseline and "All" data points consist of sampling from the entire fixed set of label functions.

### Discriminative Model Performance

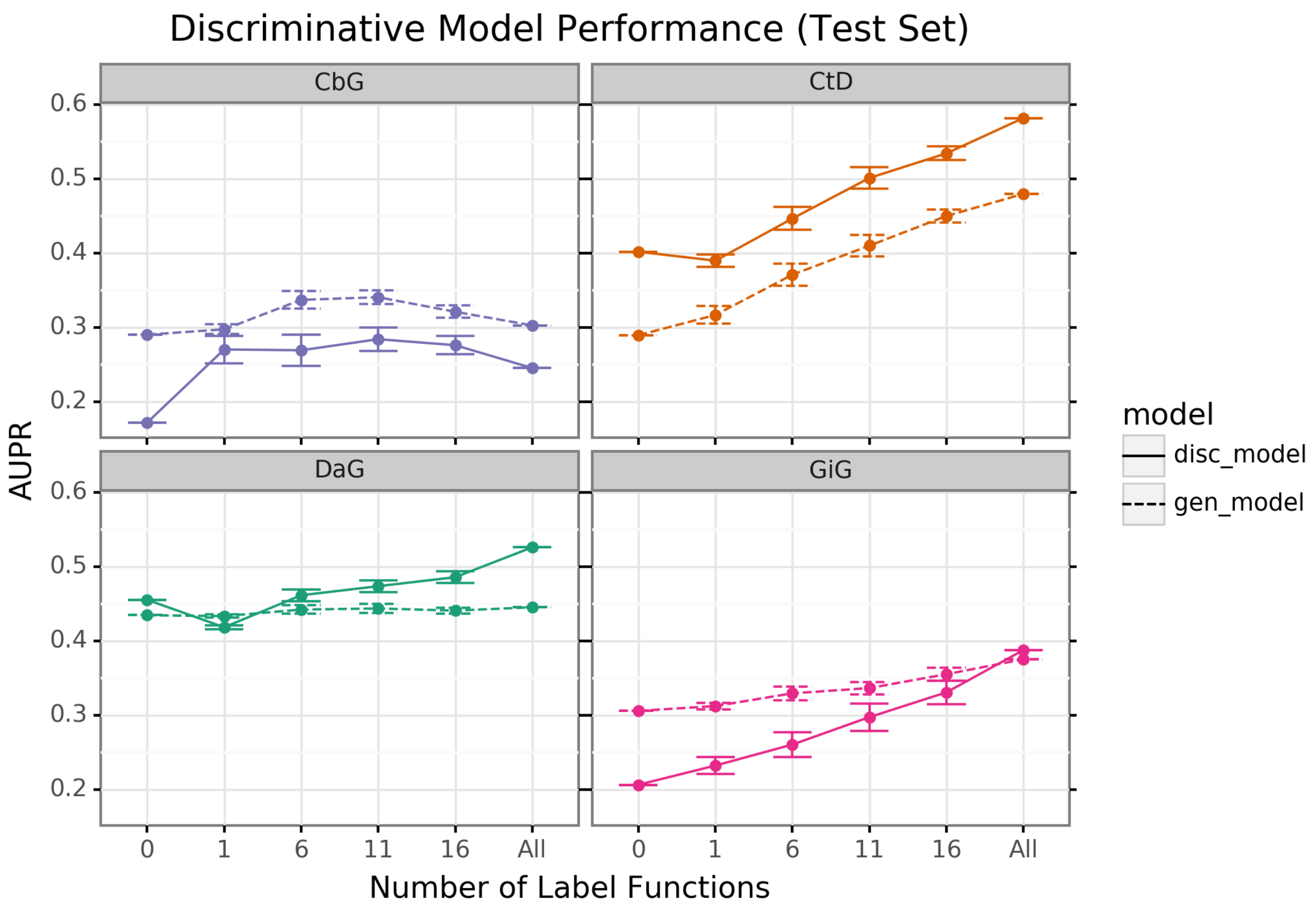


**Figure 8:** The discriminator model improves performance as the number of edge-specific label functions is added to the baseline model. The line plot headers represents the specific edge type the discriminator model is trying to predict. The x-axis shows the number of randomly sampled label functions incorporated on top of the baseline model (point at 0). The y axis shows the area under the precision recall curve (AUPR). Each datapoint shows the average of 50 sample runs, while the error bars represents the 95% confidence interval at each point. The baseline and "All" data points consist of sampling from the entire fixed set of label functions.

### Discriminative Model Calibration

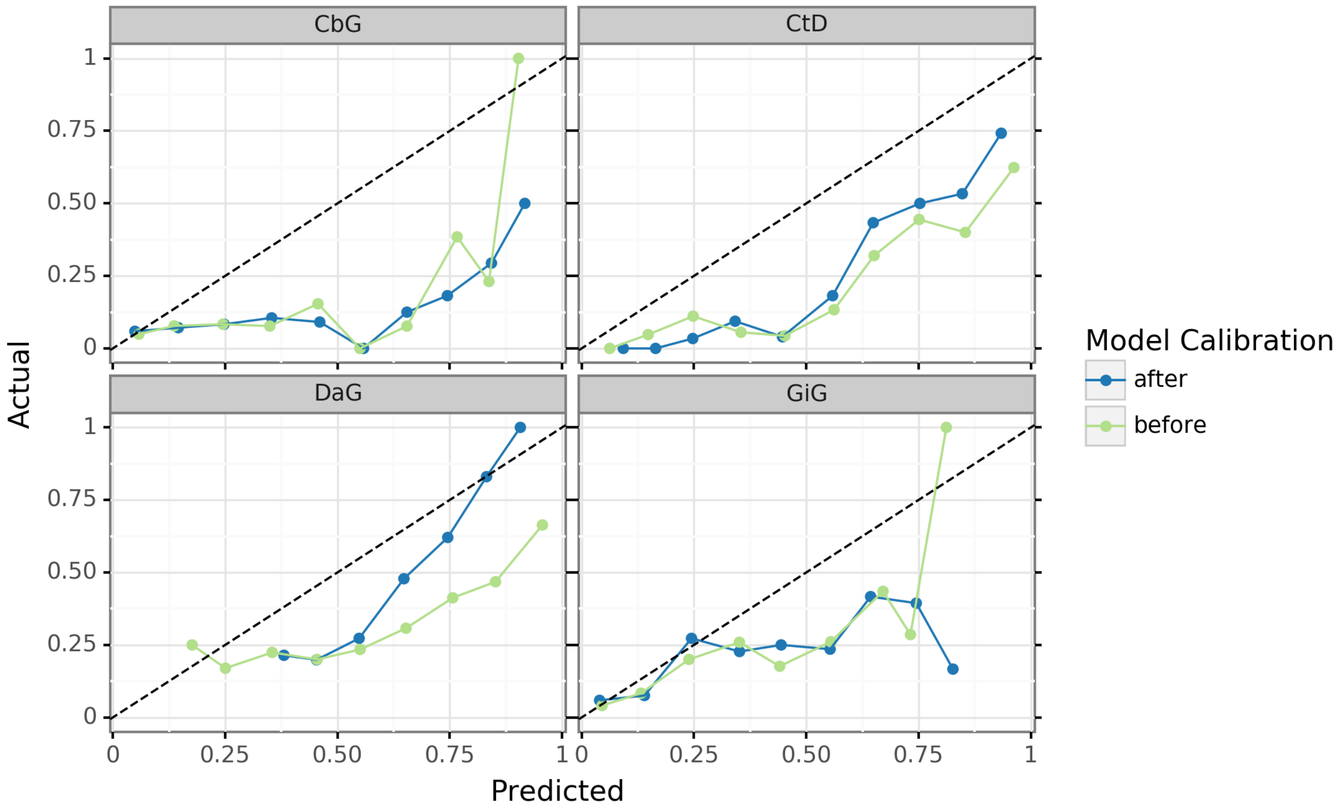


**Figure 9:** Deep learning models are overconfident in their predictions and need to be calibrated after training. These are calibration plots for the discriminative model, where the green line represents the predictions before calibration and the blue line shows predictions after calibration. Data points that lie closer to the diagonal line show better model calibration, while data points far from the diagonal line show poor performance. A perfectly calibrated model would align straight along the diagonal line.

Even deep learning models with impressive AUROC and AUPR statistics can be subject to poor calibration. Typically, these models are overconfident in their predictions [82,83]. We attempted to use temperature scaling to fix the calibration of the best performing discriminative models (Figure 9). Before calibration (green lines), our models were aligned with the ideal calibration only when predicting low probability scores (close to 0.25). Applying the temperature scaling calibration algorithm (blue lines) did not substantially improve the calibration of the model in most cases. The exception to this pattern is the Disease associates Gene (DaG) model where high confidence scores are shown to be better calibrated. Overall, calbrating deep learning models is a nontrivial task that requires more complex approaches to accomplish.

### Text Mined Edges Can Expand a Database-derived Knowledge Graph

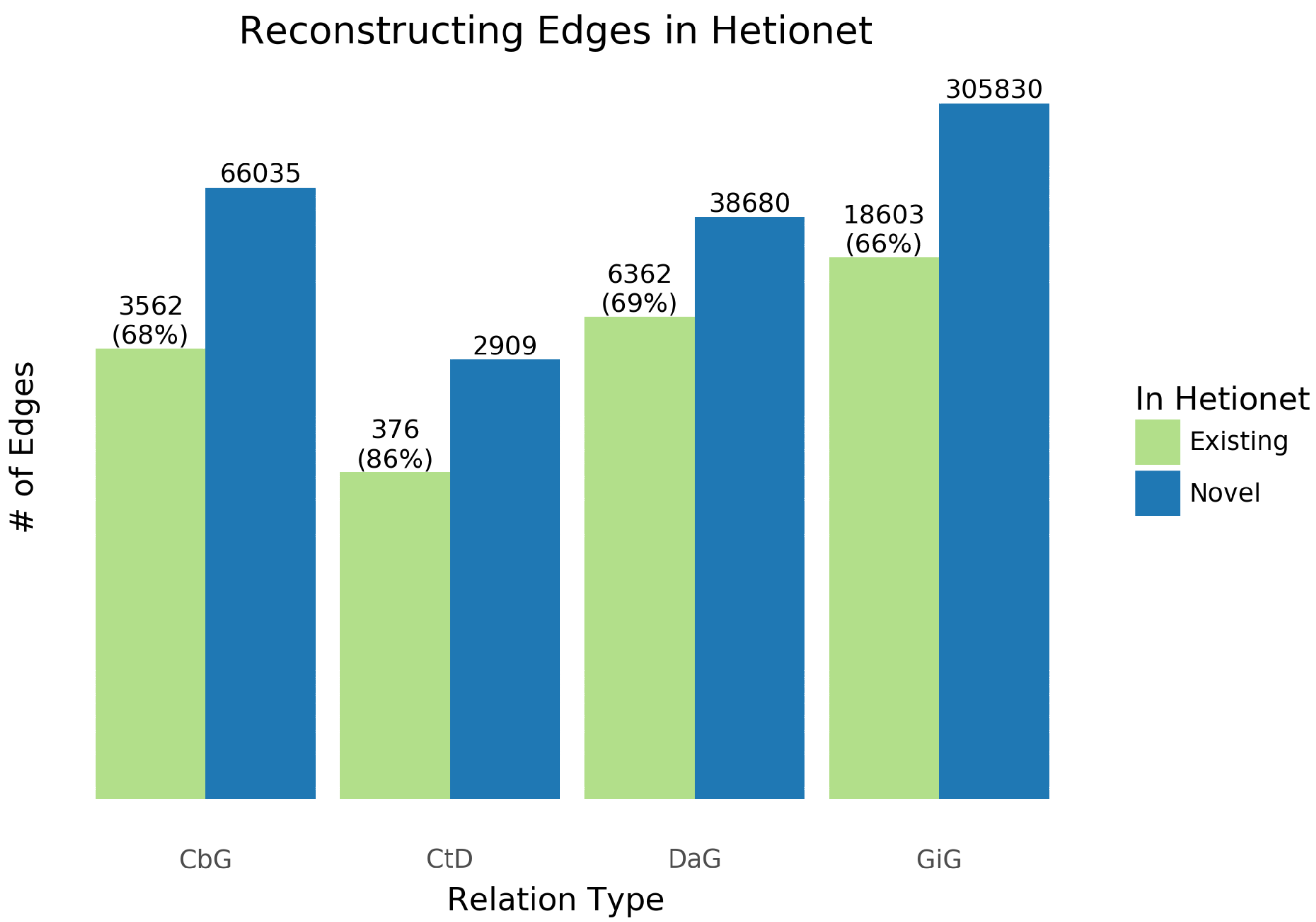


**Figure 10:** Text-mined edges recreate a substantial fraction of an existing knowledge graph and include new predictions. This bar chart shows the number of edges we can successfully recall in green and shows the number of new edges that can be added in blue.
The recall for the Hetionet v1 knowledge graph is shown as a percentage in parentheses. For example, for the Compound treats Disease (CtD) edge our method recalls 85% of existing edges and adds 6,088 new edges.

One of the goals in our work is to measure the extent to which learning multiple edge types could construct a biomedical knowledge graph. Using Hetionet v1 as an evaluation set, we measured this framework's recall and quantified how many new edges could be added with high confidence. Overall, we were able to recall more than half of preexisting edges for all edge types (Figure 10) and report our top ten scoring sentences for each edge type in Supplemental Table 11. Our best recall is with the Compound treats Disease (CtD) edge type, where we retain 85% of preexisting edges. Plus, we can add over 6,000 new edges to that category. In contrast, we could only recall close to 70% of existing edges for the other categories; however, we can add over 40,000 novel edges to each category. This highlights the fact that Hetionet v1 is missing a compelling amount of biomedical information and this framework is a viable way to close the information gap.

### Comparison with CoCoScore using Hetionet v1 as an Evaluation Set

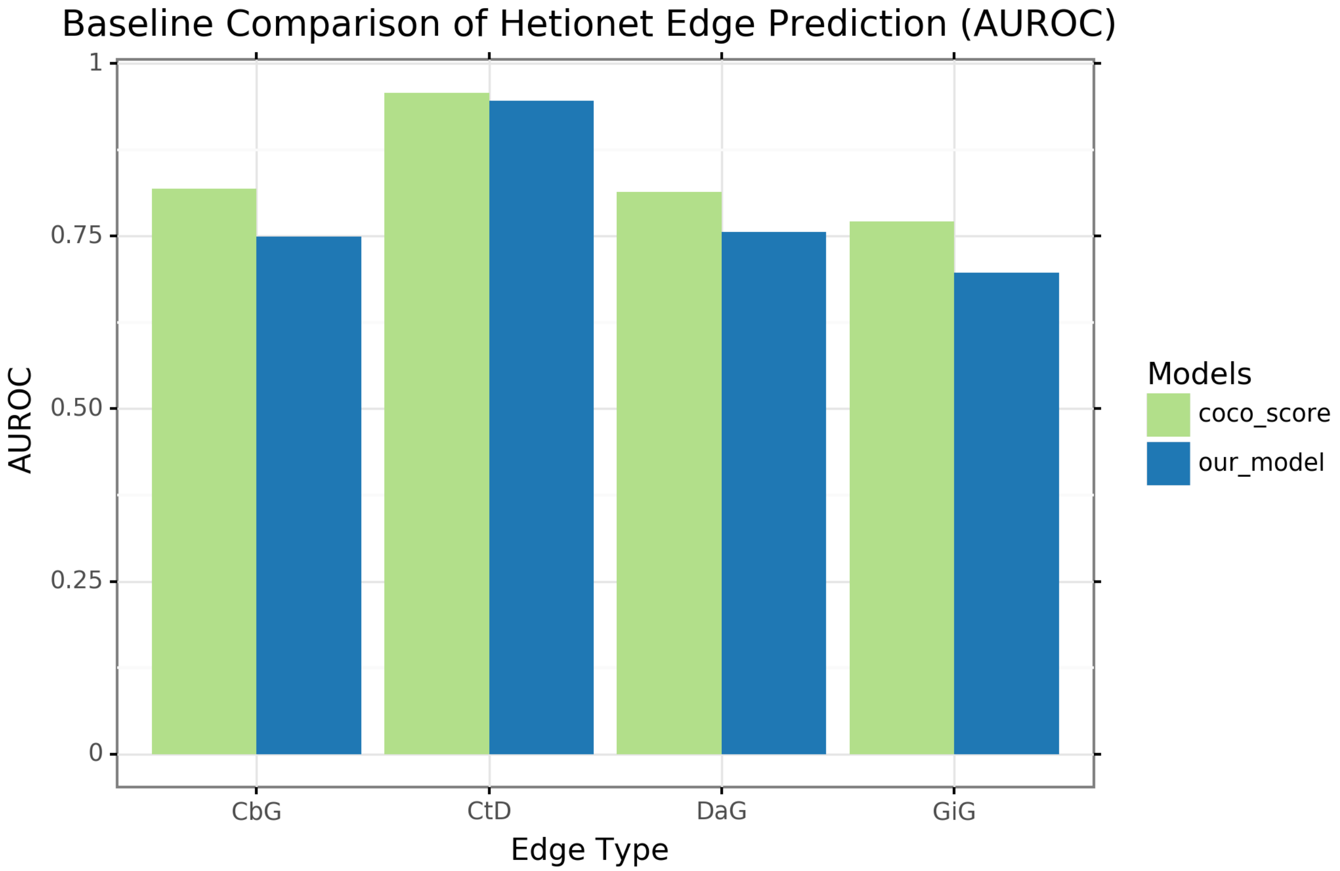


**Figure 11:** Our extractor shows similar performance to a previously published method when using Hetionet v1 as an evaluation set. We compared our model (blue) with the CoCoScore model [30] (green). The y axis represents AUROC and the x axis represents the edge type both models are trying to predict.

Our model showed promising performance in terms of recalling edges in Hetionet v1. We assessed our model's performance relative to a recently published method [30]. Though our method is primarily designed to predict assertions, not edges, we compared performance at an edge level because this was available for CoCoScore. We found that a simple summary approach, max sentence score, provided comparable performance to the CoCoScore for the compound treats disease (CtD) edge type and slightly poorer performance for other edge types (Supplemental Figure 11). Sentence-level scores can be integrated in multiple ways, and approaches that consider more complexity (e.g., the number of sentences with high-probability) should be evaluated in future work.

### Supplemental Tables

#### Distribution of Candidate Sentences

**Table 2:** Statistics of Candidate Sentences. We sorted each candidate sentence into a training, tuning and testing set. Numbers in parentheses show the number of positives and negatives that resulted from the hand-labeling process.

| Relationship | Train | Tune | Test |
|---|---|---|---|
| Disease Associates Gene | 2.35 M | 31K (397+, 603-) | 313K (351+, 649-) |
| Compound Binds Gene | 1.7M | 468K (37+, 463-) | 227k (31+, 469-) |
| Compound Treats Disease | 1.013M | 96K (96+, 404-) | 32K (112+, 388-) |
| Gene Interacts Gene | 12.6M | 1.056M (60+, 440-) | 257K (76+, 424-) |

#### Discriminative Model Calibration Tables

**Table 3:** Contains the top ten Disease-associates-Gene confidence scores before and after model calbration. Disease mentions are highlighted in brown and Gene mentions are highlighted in blue.

| Disease Name | Gene Symbol | Text | Before Calibration | After Calibration |
|---|---|---|---|---|
| prostate cancer | DKK1 | conclusion : high dkk-1 serum levels are associated with a poor survival in patients with prostate cancer . | 0.999 | 0.916 |
| breast cancer | ERBB2 | conclusion : her-2 / neu overexpression in primary breast carcinoma is correlated with patients ' age ( under age 50 ) and calcifications at mammography . | 0.998 | 0.906 |
| breast cancer | ERBB2 | the results of multiple linear regression analysis , with her2 as the dependent variable , showed that family history of breast cancer was significantly associated with elevated her2 levels in the tumors ( p = 0.0038 ) , after controlling for the effects of age , tumor estrogen receptor , and dna index . | 0.998 | 0.904 |
| colon cancer | SP3 | ba also decreased expression of sp1 , sp3 and sp4 transcription factors which are overexpressed in colon cancer cells and decreased levels of several sp-regulated genes including survivin , vascular endothelial growth factor , p65 sub-unit of nfkb , epidermal growth factor receptor , cyclin d1 , and pituitary tumor transforming gene-1 . | 0.998 | 0.902 |
| breast cancer | ERBB2 | in breast cancer , overexpression of her2 is associated with an aggressive tumor phenotype and poor prognosis . | 0.998 | 0.898 |
| breast cancer | BCL2 | in clinical breast cancer samples , high bcl2 expression was associated with poor prognosis . | 0.997 | 0.886 |
| adrenal gland cancer | TP53 | the mechanisms of adrenal tumorigenesis remain poorly established ; the r337h germline mutation in the p53 gene has previously been associated with acts in brazilian children . | 0.996 | 0.883 |
| prostate cancer | AR | the androgen receptor was expressed in all primary and metastatic prostate cancer tissues and no mutations were identified . | 0.996 | 0.881 |
| urinary bladder cancer | PIK3CA | conclusions : increased levels of fgfr3 and pik3ca mutated dna in urine and plasma are indicative of later progression and metastasis in bladder cancer . | 0.995 | 0.866 |
| ovarian cancer | EPAS1 | the log-rank test showed that nuclear positive immunostaining for hif-1alpha ( p = .002 ) and cytoplasmic positive immunostaining for hif-2alpha ( p = .0112 ) in tumor cells are associated with poor prognosis of patients with ovarian carcinoma . | 0.994 | 0.86 |

**Table 4:** Contains the bottom ten Disease-associates-Gene confidence scores before and after model calbration. Disease mentions are highlighted in brown and Gene mentions are highlighted in blue.

| Disease Name | Gene Symbol | Text | Before Calibration | After Calibration |
|---|---|---|---|---|

| Disease Name | Gene Symbol | Text | Before Calibration | After Calibration |
|---|---|---|---|---|
| endogenous depression | EP300 | from a clinical point of view , p300 amplitude should be considered as a psychophysiological index of suicidal risk in major depressive disorder . | 0.202 | 0.379 |
| Alzheimer's disease | PDK1 | from prion diseases to alzheimer 's disease : a common therapeutic target , [pdk1 ] . | 0.2 | 0.378 |
| endogenous depression | HTR1A | gepirone , a selective serotonin ( 5ht1a ) partial agonist in the treatment of major depression . | 0.199 | 0.378 |
| Gilles de la Tourette syndrome | FGF9 | there were no differences in gender distribution , age at tic onset or td diagnosis , tic severity , proportion with current diagnoses of ocd/oc behavior or attention deficit hyperactivity disorder ( adhd ) , cbcl internalizing , externalizing , or total problems scores , ygtss scores , or gaf scores . | 0.185 | 0.37 |
| hematologic cancer | MLANA | methods : the sln sections ( n = 214 ) were assessed by qrt assay for 4 established messenger rna biomarkers : mart-1 , mage-a3 , galnac-t , and pax3 . | 0.18 | 0.368 |
| endogenous depression | MAOA | alpha 2-adrenoceptor responsivity in depression : effect of chronic treatment with moclobemide , a selective mao-a-inhibitor , versus maprotiline . | 0.179 | 0.367 |
| chronic kidney failure | B2M | to evaluate comparative beta 2-m removal we studied six stable end-stage renal failure patients during high-flux 3-h haemodialysis , haemodia-filtration , and haemofiltration , using acrylonitrile , cellulose triacetate , polyamide and polysulphone capillary devices . | 0.178 | 0.366 |
| hematologic cancer | C7 | serum antibody responses to four haemophilus influenzae type b capsular polysaccharide-protein conjugate vaccines ( prp-d , hboc , c7p , and prp-t ) were studied and compared in 175 infants , 85 adults and 140 2-year-old children . | 0.174 | 0.364 |

| Disease Name | Gene Symbol | Text | Before Calibration | After Calibration |
|---|---|---|---|---|
| hypertension | AVP | portohepatic pressures , hepatic function , and blood gases in the combination of nitroglycerin and vasopressin : search for additive effects in cirrhotic portal hypertension . | 0.168 | 0.361 |
| endogenous depression | GAD1 | within-individual deflections in gad , physical , and social symptoms predicted later deflections in depressive symptoms , and deflections in depressive symptoms predicted later deflections in gad and separation anxiety symptoms . | 0.149 | 0.349 |

**Table 5:** Contains the top ten Compound-treats-Disease confidence scores after model calbration. Disease mentions are highlighted in brown and Compound mentions are highlighted in red.

| Compound Name | Disease Name | Text | Before Calibration | After Calibration |
|---|---|---|---|---|
| Prazosin | hypertension | experience with prazosin in the treatment of hypertension . | 0.997 | 0.961 |
| Methyldopa | hypertension | oxprenolol plus cyclopenthiazide-kcl versus methyldopa in the treatment of hypertension . | 0.997 | 0.961 |
| Methyldopa | hypertension | atenolol and methyldopa in the treatment of hypertension . | 0.996 | 0.957 |
| Prednisone | asthma | prednisone and beclomethasone for treatment of asthma . | 0.995 | 0.953 |
| Sulfasalazine | ulcerative colitis | sulphasalazine , used in the treatment of ulcerative colitis , is cleaved in the colon by the metabolic action of colonic bacteria on the diazo bond to release 5-aminosalicylic acid ( 5-asa ) and sulpharidine . | 0.994 | 0.949 |
| Prazosin | hypertension | letter : prazosin in treatment of hypertension . | 0.994 | 0.949 |
| Methylprednisolone | asthma | use of tao without methylprednisolone in the treatment of severe asthma . | 0.994 | 0.948 |
| Budesonide | asthma | thus , a regimen of budesonide treatment that consistently attenuates bronchial responsiveness in asthmatic subjects had no effect in these men ; larger and longer trials will be required to establish whether a subgroup of smokers shows a favorable response . | 0.994 | 0.946 |
| Methyldopa | hypertension | pressor and chronotropic responses to bilateral carotid occlusion ( bco ) and tyramine were also markedly reduced following treatment with methyldopa , which is consistent with the clinical findings that chronic methyldopa treatment in hypertensive patients impairs cardiovascular reflexes . | 0.994 | 0.946 |
| Fluphenazine | schizophrenia | low dose fluphenazine decanoate in maintenance treatment of schizophrenia . | 0.994 | 0.946 |

**Table 6:** Contains the bottom ten Compound-treats-Disease confidence scores before and after model calbration. Disease mentions are highlighted in brown and Compound mentions are highlighted in red.

| Compound Name | Disease Name | Text | Before Calibration | After Calibration |
|---|---|---|---|---|
| Indomethacin | hypertension | effects of indomethacin in rabbit renovascular hypertension . | 0.033 | 0.13 |
| Alprazolam | panic disorder | according to logistic regression analysis , the relationships between plasma alprazolam concentration and response , as reflected by number of panic attacks reported , phobia ratings , physicians ' and patients ' ratings of global improvement , and the emergence of side effects , were significant . | 0.03 | 0.124 |
| Mestranol | polycystic ovary syndrome | the binding capacity of plasma testosterone-estradiol-binding globulin ( tebg ) and testosterone ( t ) levels were measured in four women with proved polycystic ovaries and three women with a clinical diagnosis of polycystic ovarian disease before , during , and after administration of norethindrone , 2 mg. , and mestranol , 0.1 mg . | 0.03 | 0.123 |
| Creatine | coronary artery disease | during successful and uncomplicated angioplasty ( ptca ) , we studied the effect of a short lasting myocardial ischemia on plasma creatine kinase , creatine kinase mb-activity , and creatine kinase mm-isoforms ( mm1 , mm2 , mm3 ) in 23 patients . | 0.028 | 0.12 |
| Creatine | coronary artery disease | in 141 patients with acute myocardial infarction , creatine phosphokinase isoenzyme ( cpk-mb ) was determined by the activation method with dithiothreitol ( rao et al. : clin . | 0.027 | 0.117 |
| Morphine | brain cancer | the tissue to serum ratio of morphine in the hypothalamus , hippocampus , striatum , midbrain and cortex were also smaller in morphine tolerant than in non-tolerant rats . | 0.026 | 0.115 |
| Glutathione | anemia | our results suggest that an association between gsh px deficiency and hemolytic anemia need not represent a cause-and-effect relationship . | 0.026 | 0.114 |
| Dinoprostone | stomach cancer | prostaglandin e2 ( pge2 ) - and 6-keto-pgf1 alpha-like immunoactivity was measured in incubates of forestomach and gastric corpus mucosa in ( a ) unoperated rats , ( b ) rats with sham-operation of the kidneys and ( c ) rats with bilateral nephrectomy . | 0.023 | 0.107 |

| Compound Name | Disease Name | Text | Before Calibration | After Calibration |
|---|---|---|---|---|
| Creatine | coronary artery disease | the value of the electrocardiogram in assessing infarct size was studied using serial estimates of the mb isomer of creatine kinase ( ck mb ) in plasma , serial 35 lead praecordial maps in 28 patients with anterior myocardial infarction , and serial 12 lead electrocardiograms in 17 patients with inferior myocardial infarction . | 0.022 | 0.105 |
| Sulfamethazine | multiple sclerosis | quantitation and confirmation of sulfamethazine residues in swine muscle and liver by lc and gc/ms . | 0.017 | 0.093 |

**Table 7:** Contains the top ten Compound-binds-Gene confidence scores before and after model calbration. Gene mentions are highlighted in blue and Compound mentions are highlighted in red.

| Compound Name | Gene Symbol | Text | Before Calibration | After Calibration |
|---|---|---|---|---|
| Cyclic Adenosine Monophosphate | B3GNT2 | in sk-n-mc human neuroblastoma cells , the camp response to 10 nm isoproterenol ( iso ) is mediated primarily by beta 1-adrenergic receptors . | 0.903 | 0.93 |
| Indomethacin | AGT | indomethacin , a potent inhibitor of prostaglandin synthesis , is known to increase the maternal blood pressure response to angiotensin ii infusion . | 0.894 | 0.922 |
| Tretinoin | RXRA | the vitamin a derivative retinoic acid exerts its effects on transcription through two distinct classes of nuclear receptors , the retinoic acid receptor ( rar ) and the retinoid x receptor ( rxr ) . | 0.882 | 0.912 |
| Tretinoin | RXRA | the vitamin a derivative retinoic acid exerts its effects on transcription through two distinct classes of nuclear receptors , the retinoic acid receptor ( rar ) and the retinoid x receptor ( rxr ) . | 0.872 | 0.903 |

| Compound Name | Gene Symbol | Text | Before Calibration | After Calibration |
|---|---|---|---|---|
| D-Tyrosine | CSF1 | however , the extent of gap tyrosine phosphorylation induced by csf-1 was approximately 10 % of that induced by pdgf-bb in the nih3t3 fibroblasts . | 0.851 | 0.883 |
| D-Glutamic Acid | GLB1 | thus , the negatively charged side chain of glu-461 is important for divalent cation binding to beta-galactosidase . | 0.849 | 0.882 |
| D-Tyrosine | CD4 | second , we use the same system to provide evidence that the physical association of cd4 with the tcr is required for effective tyrosine phosphorylation of the tcr zeta-chain subunit , presumably reflecting delivery of p56lck ( lck ) to the tcr . | 0.825 | 0.859 |
| Calcium Chloride | TNC | the possibility that the enhanced length dependence of ca2 + sensitivity after cardiac tnc reconstitution was attributable to reduced tnc binding was excluded when the length dependence of partially extracted fast fibres was reduced to one-half the normal value after a 50 % deletion of the native tnc . | 0.821 | 0.855 |
| Metoprolol | KCNMB2 | studies in difi cells of the displacement of specific 125i-cyp binding by nonselective ( propranolol ) , beta 1-selective ( metoprolol and atenolol ) , and beta 2-selective ( ici 118-551 ) antagonists revealed only a single class of beta 2-adrenergic receptors . | 0.82 | 0.854 |

| Compound Name | Gene Symbol | Text | Before Calibration | After Calibration |
|---|---|---|---|---|
| D-Tyrosine | PLCG1 | epidermal growth factor ( egf ) or platelet-derived growth factor binding to their receptor on fibroblasts induces tyrosine phosphorylation of plc gamma 1 and stable association of plc gamma 1 with the receptor protein tyrosine kinase . | 0.818 | 0.851 |

**Table 8:** Contains the bottom ten Compound-binds-Gene confidence scores before and after model calbration. Gene mentions are highlighted in blue and Compound mentions are highlighted in red.

| Compound Name | Gene Symbol | Text | Before Calibration | After Calibration |
|---|---|---|---|---|
| Deferoxamine | TF | the mechanisms of fe uptake have been characterised using 59fe complexes of citrate , nitrilotriacetate , desferrioxamine , and 59fe added to eagle 's minimum essential medium ( mem ) and compared with human transferrin ( tf ) labelled with 59fe and iodine-125 . | 0.02 | 0.011 |
| Hydrocortisone | GH1 | group iv patients had normal basal levels of lh and normal lh , gh and cortisol responses . | 0.02 | 0.011 |
| Carbachol | INS | at the same concentration , however , iapp significantly ( p less than 0.05 ) inhibited carbachol-stimulated ( 10 ( -7 ) m ) release of insulin by 30 % , and cgrp significantly inhibited carbachol-stimulated release of insulin by 33 % when compared with the control group . | 0.02 | 0.011 |
| Adenosine | ME2 | at physiological concentrations , atp , adp , and amp all inhibit the enzyme from atriplex spongiosa and panicum miliaceum ( nad-me-type plants ) , with atp the most inhibitory species . | 0.019 | 0.01 |
| Naloxone | POMC | specifically , opioids , including 2-n-pentyloxy-2-phenyl-4-methyl-morpholine , naloxone , and beta-endorphin , have been shown to interact with il-2 receptors ( 134 ) and regulate production of il-1 and il-2 ( 48-50 , 135 ) . | 0.018 | 0.01 |
| Cortisone acetate | POMC | sarcoidosis therapy with cortisone and acth – the role of acth therapy . | 0.017 | 0.009 |
| Epinephrine | INS | thermogenic effect of thyroid hormones : interactions with epinephrine and insulin . | 0.017 | 0.009 |
| Aldosterone | KNG1 | important vasoconstrictor , fluid - and sodium-retaining factors are the renin-angiotensin-aldosterone system , sympathetic nerve activity , and vasopressin ; vasodilator , volume , and sodium-eliminating factors are atrial natriuretic peptide , vasodilator prostaglandins like prostacyclin and prostaglandin e2 , dopamine , bradykinin , and possibly , endothelial derived relaxing factor ( edrf ) . | 0.016 | 0.008 |
| D-Leucine | POMC | cross-reactivities of leucine-enkephalin and beta-endorphin with the eia were less than 0.1 % , while that with gly-gly-phe-met and oxidized gly-gly-phe-met were 2.5 % and 10.2 % , respectively . | 0.011 | 0.005 |

| Compound Name | Gene Symbol | Text | Before Calibration | After Calibration |
|---|---|---|---|---|
| Estriol | LGALS1 | [ diagnostic value of serial determination of estriol and hpl in plasma and of total estrogens in 24-h-urine compared to single values for diagnosis of fetal danger ] . | 0.01 | 0.005 |

**Table 9:** Contains the top ten Gene-interacts-Gene confidence scores before and after model calbration. Both gene mentions highlighted in blue.

| Gene1 Symbol | Gene2 Symbol | Text | Before Calibration | After Calibration |
|---|---|---|---|---|
| ESR1 | HSP90AA1 | previous studies have suggested that the 90-kda heat shock protein ( hsp90 ) interacts with the er , thus stabilizing the receptor in an inactive state . | 0.812 | 0.864 |
| TP53 | TP73 | cyclin g interacts with p53 as well as p73 , and its binding to p53 or p73 presumably mediates downregulation of p53 and p73 . | 0.785 | 0.837 |
| TP53 | AKT1 | treatment of c81 cells with ly294002 resulted in an increase in the p53-responsive gene mdm2 , suggesting a role for akt in the tax-mediated regulation of p53 transcriptional activity . | 0.773 | 0.825 |
| ABCB1 | NR1I3 | valproic acid induces cyp3a4 and mdr1 gene expression by activation of constitutive androstane receptor and pregnane x receptor pathways . | 0.762 | 0.813 |

| Gene1 Symbol | Gene2 Symbol | Text | Before Calibration | After Calibration |
|---|---|---|---|---|
| PTH2R | PTH2 | thus , the juxtamembrane receptor domain specifies the signaling and binding selectivity of tip39 for the pth2 receptor over the pth1 receptor . | 0.761 | 0.812 |
| CCND1 | ABL1 | synergy with v-abl depended on a motif in cyclin d1 that mediates its binding to the retinoblastoma protein , suggesting that abl oncogenes in part mediate their mitogenic effects via a retinoblastoma protein-dependent pathway . | 0.757 | 0.808 |
| CTNND1 | CDH1 | these complexes are formed independently of ddr1 activation and of beta-catenin and p120-catenin binding to e-cadherin ; they are ubiquitous in epithelial cells . | 0.748 | 0.798 |
| CSF1 | CSF1R | this is in agreement with current thought that the c-fms proto-oncogene product functions as the csf-1 receptor specific to this pathway . | 0.745 | 0.795 |

| Gene1 Symbol | Gene2 Symbol | Text | Before Calibration | After Calibration |
|---|---|---|---|---|
| EZR | CFTR | without ezrin binding , the cytoplasmic tail of cftr only interacts strongly with the first amino-terminal pdz domain to form a 1:1 c-cftr . | 0.732 | 0.78 |
| SRC | PIK3CG | we have demonstrated that the sh2 ( src homology 2 ) domains of the 85 kda subunit of pi-3k are sufficient to mediate binding of the pi-3k complex to tyrosine phosphorylated , but not non-phosphorylated il-2r beta , suggesting that tyrosine phosphorylation is an integral component of the activation of pi-3k by the il-2r . | 0.731 | 0.78 |

**Table 10:** Contains the bottom ten Gene-interacts-Gene confidence scores before and after model calbration. Both gene mentions highlighted in blue.

| Gene1 Symbol | Gene2 Symbol | Text | Before Calibration | After Calibration |
|---|---|---|---|---|
| AGTR1 | ACE | result ( s ) : the luteal tissue is the major site of ang ii , ace , at1r , and vegf , with highest staining intensity found during the midluteal phase and at pregnancy . | 0.009 | 0.003 |

| Gene1 Symbol | Gene2 Symbol | Text | Before Calibration | After Calibration |
|---|---|---|---|---|
| ABCE1 | ABCF2 | in relation to normal melanocytes , abcb3 , abcb6 , abcc2 , abcc4 , abce1 and abcf2 were significantly increased in melanoma cell lines , whereas abca7 , abca12 , abcb2 , abcb4 , abcb5 and abcd1 showed lower expression levels . | 0.008 | 0.002 |
| IL4 | IFNG | in contrast , il-13ralpha2 mrna expression was up-regulated by ifn-gamma plus il-4 . | 0.007 | 0.002 |
| FCAR | CD79A | we report here the presence of circulating soluble fcalphar ( cd89 ) - iga complexes in patients with igan . | 0.007 | 0.002 |
| IL4 | VCAM1 | similarly , il-4 induced vcam-1 expression and augmented tnf-alpha-induced expression on huvec but did not affect vcam-1 expression on hdmec . | 0.007 | 0.002 |

| Gene1 Symbol | Gene2 Symbol | Text | Before Calibration | After Calibration |
| --- | --- | --- | --- | --- |
| IL2 | IFNG | prostaglandin e2 at priming of naive cd4 + t cells inhibits acquisition of ability to produce ifn-gamma and il-2 , but not il-4 and il-5 . | 0.006 | 0.002 |
| IL2 | FOXP3 | il-1b promotes tgf-b1 and il-2 dependent foxp3 expression in regulatory t cells . | 0.006 | 0.002 |
| IL2 | IFNG | the detailed distribution of lymphokine-producing cells showed that il-2 and ifn-gamma-producing cells were located mainly in the follicular areas . | 0.005 | 0.001 |
| IFNG | IL10 | results : we found weak mrna expression of interleukin-4 ( il-4 ) and il-5 , and strong expression of il-6 , il-10 and ifn-gamma before therapy . | 0.005 | 0.001 |
| PIK3R1 | PTEN | both pten ( pi3k antagonist ) and pp2 ( unspecific phosphatase ) were down-regulated . | 0.005 | 0.001 |

### Top Ten Sentences for Each Edge Type

**Table 11:** Contains the top ten predictions for each edge type. Highlighted words represent entities mentioned within the given sentence.

| Edge Type | Source Node | Target Node | Generative Model Prediction | Discriminative Model Prediction | Number of Sentences | In Hetionet | Text |
|---|---|---|---|---|---|---|---|
| DaG | urinary bladder cancer | TP53 | 1 | 0.945 | 2112 | Existing | conclusion : our findings indicate that the dsp53-285 can upregulate wild-type p53 expression in human bladder cancer cells through rna activation , and suppresses cells proliferation and metastasis in vitro and in vivo . |
| DaG | ovarian cancer | EGFR | 1 | 0.937 | 1330 | Existing | conclusion : our data showed that increased expression of egfr is associated with poor prognosis of patients with eoc and dacomitinib may act as a novel , useful chemotherapy drug . |
| DaG | stomach cancer | TP53 | 1 | 0.937 | 2679 | Existing | conclusion : this meta-analysis suggests that p53 arg72pro polymorphism is associated with increased risk of gastric cancer in asians . |
| DaG | lung cancer | TP53 | 1 | 0.936 | 6813 | Existing | conclusion : these results suggest that high expression of the p53 oncoprotein is a favorable prognostic factor in a subset of patients with nsclc . |
| DaG | breast cancer | TCF7L2 | 1 | 0.936 | 56 | Existing | this meta-analysis demonstrated that tcf7l2 gene polymorphisms ( rs12255372 and rs7903146 ) are associated with an increased susceptibility to breast cancer . |
| DaG | skin cancer | COX2 | 1 | 0.935 | 73 | Novel | elevated expression of cox-2 has been associated with tumor progression in skin cancer through multiple mechanisms . |

| Edge Type | Source Node | Target Node | Generative Model Prediction | Discriminative Model Prediction | Number of Sentences | In Hetionet | Text |
|---|---|---|---|---|---|---|---|
| DaG | thyroid cancer | VEGFA | 1 | 0.933 | 592 | Novel | as a conclusion , we suggest that vegf g +405 c polymorphism is associated with increased risk of ptc . |
| DaG | stomach cancer | EGFR | 1 | 0.933 | 1237 | Existing | recently , high lymph node ratio is closely associated with egfr expression in advanced gastric cancer . |
| DaG | liver cancer | GPC3 | 1 | 0.933 | 1944 | Novel | conclusions serum gpc3 was overexpressed in hcc patients . |
| DaG | stomach cancer | CCR6 | 1 | 0.931 | 24 | Novel | the cox regression analysis showed that high expression of ccr6 was an independent prognostic factor for gc patients . |
| CtD | Sorafenib | liver cancer | 1 | 0.99 | 6672 | Existing | tace plus sorafenib for the treatment of hepatocellular carcinoma : final results of the multicenter socrates trial . |
| CtD | Methotrexate | rheumatoid arthritis | 1 | 0.989 | 14546 | Existing | comparison of low-dose oral pulse methotrexate and placebo in the treatment of rheumatoid arthritis . |
| CtD | Auranofin | rheumatoid arthritis | 1 | 0.988 | 419 | Existing | auranofin versus placebo in the treatment of rheumatoid arthritis . |
| CtD | Lamivudine | hepatitis B | 1 | 0.988 | 6709 | Existing | randomized controlled trials ( rcts ) comparing etv with lam for the treatment of hepatitis b decompensated cirrhosis were included . |

| Edge Type | Source Node | Target Node | Generative Model Prediction | Discriminative Model Prediction | Number of Sentences | In Hetionet | Text |
|---|---|---|---|---|---|---|---|
| CtD | Doxorubicin | urinary bladder cancer | 1 | 0.988 | 930 | Existing | 17-year follow-up of a randomized prospective controlled trial of adjuvant intravesical doxorubicin in the treatment of superficial bladder cancer . |
| CtD | Docetaxel | breast cancer | 1 | 0.987 | 5206 | Existing | currently , randomized phase iii trials have demonstrated that docetaxel is an effective strategy in the adjuvant treatment of breast cancer . |
| CtD | Cimetidine | psoriasis | 0.999 | 0.987 | 12 | Novel | cimetidine versus placebo in the treatment of psoriasis . |
| CtD | Olanzapine | schizophrenia | 1 | 0.987 | 3324 | Novel | a double-blind , randomised comparative trial of amisulpride versus olanzapine in the treatment of schizophrenia : short-term results at two months . |
| CtD | Fulvestrant | breast cancer | 1 | 0.987 | 826 | Existing | phase iii clinical trials have demonstrated the clinical benefit of fulvestrant in the endocrine treatment of breast cancer . |
| CtD | Pimecrolimus | atopic dermatitis | 1 | 0.987 | 531 | Existing | introduction : although several controlled clinical trials have demonstrated the efficacy and good tolerability of 1 % pimecrolimus cream for the treatment of atopic dermatitis , the results of these trials may not apply to real-life usage . |

| Edge Type | Source Node | Target Node | Generative Model Prediction | Discriminative Model Prediction | Number of Sentences | In Hetionet | Text |
|---|---|---|---|---|---|---|---|
| CbG | Gefitinib | EGFR | 1 | 0.99 | 8746 | Existing | morphologic features of adenocarcinoma of the lung predictive of response to the epidermal growth factor receptor kinase inhibitors erlotinib and gefitinib . |
| CbG | Adenosine | EGFR | 1 | 0.987 | 644 | Novel | it is well established that inhibiting atp binding within the egfr kinase domain regulates its function . |
| CbG | Rosiglitazone | PPARG | 1 | 0.987 | 1498 | Existing | rosiglitazone is a potent peroxisome proliferator-activated receptor gamma agonist that decreases hyperglycemia by reducing insulin resistance in patients with type 2 diabetes mellitus . |
| CbG | D-Tyrosine | INSR | 0.998 | 0.987 | 1713 | Novel | this result suggests that tyrosine phosphorylation of phosphatidylinositol 3-kinase by the insulin receptor kinase may increase the specific activity of the former enzyme in vivo . |
| CbG | D-Tyrosine | IGF1 | 0.998 | 0.983 | 819 | Novel | affinity-purified insulin-like growth factor i receptor kinase is activated by tyrosine phosphorylation of its beta subunit . |
| CbG | Pindolol | HTR1A | 1 | 0.983 | 175 | Existing | pindolol , a betablocker with weak partial 5-ht1a receptor agonist activity has been shown to produce a more rapid onset of antidepressant action of ssris . |

| Edge Type | Source Node | Target Node | Generative Model Prediction | Discriminative Model Prediction | Number of Sentences | In Hetionet | Text |
|---|---|---|---|---|---|---|---|
| CbG | Progesterone | SHBG | 1 | 0.981 | 492 | Existing | however , dng also elicits properties of progesterone derivatives like neutrality in metabolic and cardiovascular system and considerable antiandrogenic activity , the latter increased by lack of binding to shbg as specific property of dng . |
| CbG | Mifepristone | AR | 1 | 0.98 | 78 | Existing | ru486 bound to the androgen receptor . |
| CbG | Alfentanil | OPRM1 | 1 | 0.979 | 10 | Existing | purpose : alfentanil is a high potency mu opiate receptor agonist commonly used during presurgical induction of anesthesia . |
| CbG | Candesartan | AGTR1 | 1 | 0.979 | 36 | Existing | tcv-116 is a new , nonpeptide , angiotensin ii type-1 receptor antagonist that acts as a specific inhibitor of the renin-angiotensin system . |
| GiG | BRCA2 | BRCA1 | 0.972 | 0.984 | 12257 | Novel | a total of 9 families ( 16 % ) showed mutations in the brca1 gene , including the one new mutation identified in this study ( 5382insc ) , and 12 families ( 21 % ) presented mutations in the brca2 gene . |
| GiG | MDM2 | TP53 | 0.938 | 0.978 | 17128 | Existing | no mutations in the tp53 gene have been found in samples with amplification of mdm2 . |

| Edge Type | Source Node | Target Node | Generative Model Prediction | Discriminative Model Prediction | Number of Sentences | In Hetionet | Text |
|---|---|---|---|---|---|---|---|
| GiG | BRCA1 | BRCA2 | 1 | 0.978 | 12257 | Existing | pathogenic truncating mutations in the brca1 gene were found in two tumor samples with allelic losses , whereas no mutations were identified in the brca2 gene . |
| GiG | KRAS | TP53 | 0.992 | 0.971 | 4106 | Novel | mutations in the p53 gene did not correlate with mutations in the c-k-ras gene , indicating that colorectal cancer can develop through pathways independent not only of the presence of mutations in any of these genes but also of their cooperation . |
| GiG | TP53 | HRAS | 0.992 | 0.969 | 451 | Novel | pathologic examination of the uc specimens from aa-exposed patients identified heterozygous hras changes in 3 cases , and deletion or replacement mutations in the tp53 gene in 4 . |
| GiG | REN | NR1H3 | 0.998 | 0.966 | 8 | Novel | nuclear receptor lxralpha is involved in camp-mediated human renin gene expression . |
| GiG | ESR2 | CYP19A1 | 0.999 | 0.96 | 159 | Novel | dna methylation , histone modifications , and binding of estrogen receptor , erb to regulatory dna sequences of cyp19a1 gene were evaluated by chromatin immunoprecipitation ( chip ) assay . |

| Edge Type | Source Node | Target Node | Generative Model Prediction | Discriminative Model Prediction | Number of Sentences | In Hetionet | Text |
|---|---|---|---|---|---|---|---|
| GiG | RET | EDNRB | 0.816 | 0.96 | 136 | Novel | mutations in the ret gene , which codes for a receptor tyrosine kinase , and in ednrb which codes for the endothelin-b receptor , have been shown to be associated with hscr in humans . |
| GiG | PKD1 | PKD2 | 1 | 0.959 | 1614 | Existing | approximately 85 % of adpkd cases are caused by mutations in the pkd1 gene , while mutations in the pkd2 gene account for the remaining 15 % of cases . |
| GiG | LYZ | CTCF | 0.999 | 0.959 | 2 | Novel | in conjunction with the thyroid receptor ( tr ) , ctcf binding to the lysozyme gene transcriptional silencer mediates the thyroid hormone response element ( tre ) - dependent transcriptional repression . |
